# Multimodal Benchmarking of Foundation Model Representations for Cellular Perturbation Response Prediction

**DOI:** 10.1101/2025.06.26.661186

**Authors:** Euxhen Hasanaj, Elijah Cole, Shahin Mohammadi, Sohan Addagudi, Xingyi Zhang, Le Song, Eric P. Xing

**Affiliations:** GenBio AI; Carnegie Mellon University; Mohamed bin Zayed University of Artificial Intelligence

## Abstract

The decreasing cost of single-cell RNA sequencing (scRNA-seq) has enabled the collection of massive scRNA-seq datasets, which are now being used to train transformer-based cell foundation models (FMs). One of the most promising applications of these FMs is perturbation response modeling. This task aims to forecast how cells will respond to drugs or genetic interventions. Accurate perturbation response models could drastically accelerate drug discovery by reducing the space of interventions that need to be tested in the wet lab. However, recent studies have shown that FM-based models often struggle to outperform simpler baselines for perturbation response prediction. A key obstacle is the lack of understanding of the components driving performance in FM-based perturbation response models. In this work, we conduct the first systematic pan-modal study of perturbation embeddings, with an emphasis on those derived from biological FMs. We benchmark their predictive accuracy, analyze patterns in their predictions, and identify the most successful representation learning strategies. Our findings offer insights into what FMs are learning and provide practical guidance for improving perturbation response modeling.

## 1. Introduction

Recent advancements in scRNA-seq technology have made it relatively inexpensive to profile the gene expression of vast numbers of individual cells (Svensson et al., 2018). It is now possible to profile 100M cells in a matter of weeks (Zhang et al., 2025). As the available data has grown, so has interest in training large foundation models (FMs) using scRNAseq data. One of the most exciting capabilities these FMs may unlock is highly accurate *perturbation response models*, which would allow us to simulate how cells would respond to drugs or genetic interventions (Gavriilidis et al., 2024). Such models would allow computational searches for promising therapies, dramatically reducing the time and expense required for wet lab experiments (Bunne et al., 2024).

So far, accurate FM-driven perturbation models have proved elusive. Several studies have found that cell FMs perform no better than much simpler models for perturbation response prediction (Ahlmann-Eltze et al., 2024; Kernfeld et al., 2023). The reasons for this performance gap are not well understood. It is unknown whether performance is limited by the perturbation embeddings, the models built on top of them, dataset size, or other factors. For instance, few prior works have systematically studied the contributions of the perturbation embeddings separately from the perturbation response model itself. In this work, we fix the perturbation response model and study perturbation embeddings from a diverse collection of modalities and representation learning strategies. This reveals new insights into what different FMs are learning and which representation learning methods are most useful for perturbation response prediction.

Our main contributions are: (i) We evaluate the predictive utility of perturbation embeddings separately from downstream perturbation response models, removing confounders and revealing which embedding representations are most useful. (ii) We conduct the first comprehensive assessment of perturbation embeddings from different modalities and models: expression-based FMs, protein FMs, DNA FMs, and embeddings based on structured biological knowledge. (iii) We introduce a novel, biologically interpretable formulation of perturbation response modeling based on predicting functional annotations (GO terms) of each perturbation.

## 2. Related Work

There is considerable interest in the idea of using biological FMs to simulate cellular behavior (Bunne et al., 2024; Song et al., 2024). There are several works that focus on evaluating FMs in the context of perturbation response modeling (Kernfeld et al., 2023; Li et al., 2024b; Ahlmann-Eltze et al., 2024; Li et al., 2024a; Csendes et al., 2025). However, when using the FMs, all of these studies use bespoke, end-to-end approaches that entangle perturbation representation with downstream prediction. This coupling introduces substantial complexity and confounding, making it difficult to isolate whether performance gains stem from superior perturbation representations or other model use protocols. Some prior works have used a fixed predictive model (Chen & Zou, 2024a; Wenteler et al., 2024), but they considered much narrower collections of FMs.

This work focuses on evaluating the utility of different perturbation embeddings. We explicitly decouple perturbation representation from perturbation response prediction. This clean separation enables rigorous, controlled evaluation of the utility of gene embeddings generated by each FM. Furthermore, our benchmarks cover a more diverse set of models and modalities, offering a more comprehensive assessment of the perturbation representation landscape. There are many specialized perturbation response models one could use on top of such embeddings (e.g. Roohani et al. (2024), Lotfollahi et al. (2023), Piran et al. (2024)), but those models are beyond the scope of our work, which intentionally fixes a single model with a consistent training protocol. Other works focus on benchmarking different classes of perturbation response models, such as Wu et al. (2024); Velez-Arce et al. (2024).

## 3. Problem Formulation

Suppose we are given a collection of scRNA-seq profiles from a perturbation experiment in which *K* different perturbations *P*_1_, …, *P*_*K*_ are tested. All profiles are normalized as described in Appendix B. If *G∈* ℕ is the number of genes we measure, then our data consists of *N* control cells *X*^*C*^ ∈ ℝ^*N×G*^ and *M*_*k*_ perturbed cells 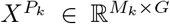 where 1 *≤ k ≤ K*. Denote the mean post-perturbation expression as 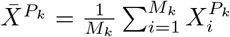 where 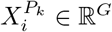 is the *i*th row of 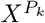. Similarly, let 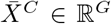 denote the mean control expression.

Our first task is to predict how a perturbation alters gene expression, by regressing from its embedding to the average change in gene expression relative to control. We call this the *pseudobulk residual expression prediction* formulation of perturbation response modeling. Formally, suppose we have an embedding *E*_*k*_ ∈ ℝ^*p*^ for each perturbation *P*_*k*_. (In this work we focus on genetic perturbations, so *E*_*k*_ would correspond to the gene being targeted.) Then the goal is to learn a model *f*_*θ*_ : ℝ^*p*^ *→* ℝ^*G*^ such that

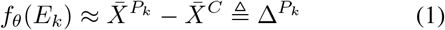

where 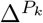 is the mean residual expression relative to control. This work always considers one cell type at a time, so our formulation does not include a parameterization of cell type. See Figs. 1, 2, 3 for results for this formulation.

**Figure 1.**
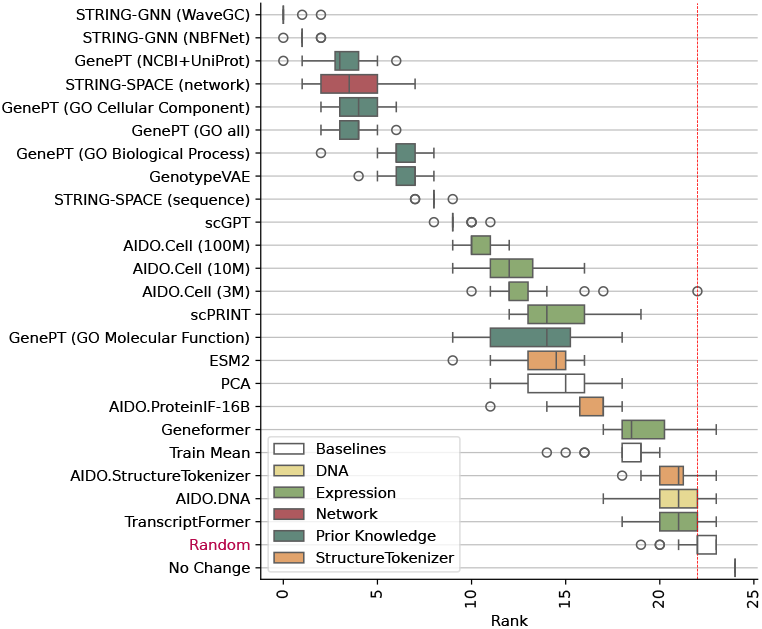
Distribution of embedding ranks in our benchmarks over 20 different trained models (4 cell lines, 5-fold cross-validation). Lower is better. Bars are colored by the type of source information used to learn the embeddings. See Fig. 2 for per-cell-line results.

**Figure 2.**
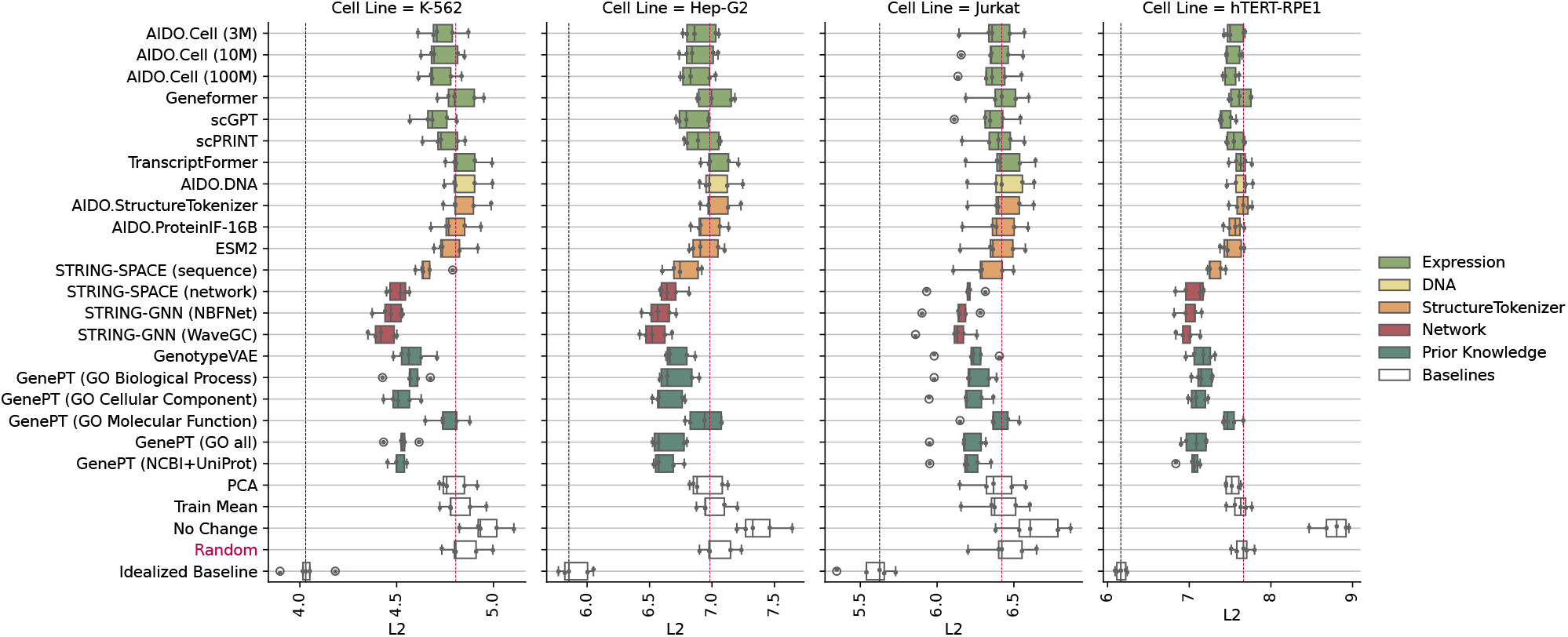
5-fold cross-validated prediction error for perturbation embeddings in four cell lines. All results are for kNN regression models with identical training and hyperparameter tuning protocols. The red dashed line corresponds to the performance of random embeddings. Bars are colored by the type of source information used to learn the embeddings.

**Figure 3.**
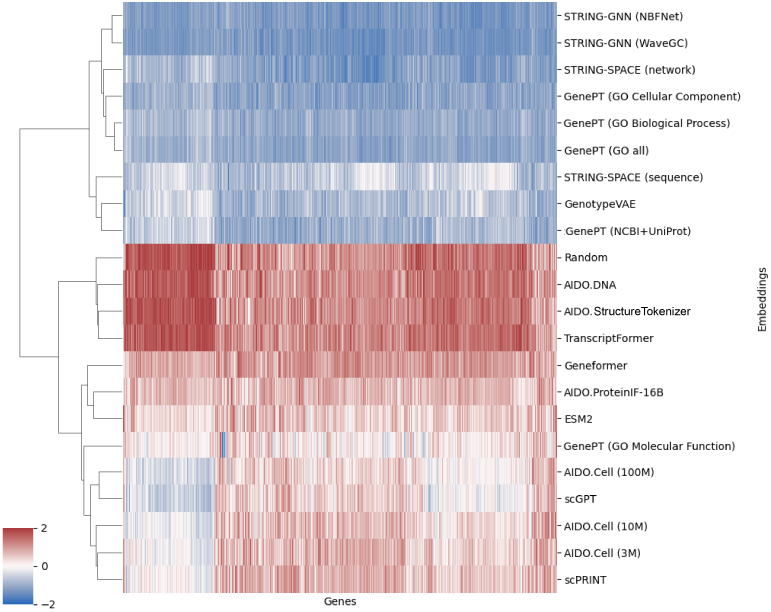
Clustermap of per-gene *ℓ*^2^ error on the K562 cell line for all embeddings. Values are *z*-scored within each gene (column), indicating whether the gene is predicted relatively well (blue) or poorly (red) compared to other methods. Embeddings with similar gene-wise error profiles will cluster together even if their overall performance differs. Color scale is clipped to [-2, 2] for contrast.

We also introduce a complementary formulation of perturbation response prediction which, to the best of our knowledge, has not been studied before: *function prediction*. Instead of predicting post-perturbation gene expression values, we (i) compute differentially expressed (DE) genes for each perturbation and (ii) perform gene set enrichment analysis (GSEA) against the gene ontology (GO) terms. For perturbation *P*_*k*_, this yields a binary vector *y*_*k*_ *∈ {*0, 1 *}*^*V*^ which indicates the presence or absence of *V* GO terms. We then train a multi-label linear classifier

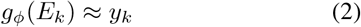

which optimizes a class-balanced binary cross-entropy loss. Results for this formulation are presented in Fig. 4.

**Figure 4.**
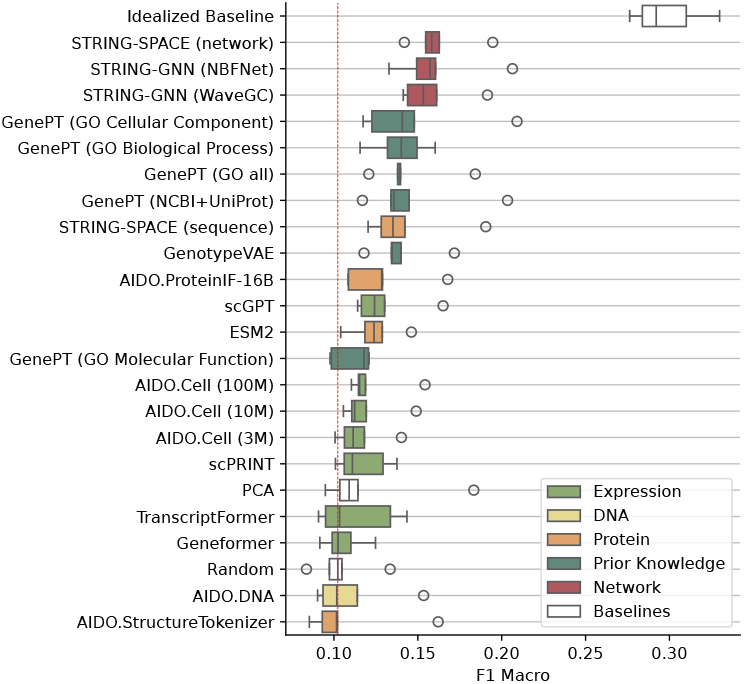
Function prediction results for the K562 cell line, sorted by average performance. Bars are colored by the type of source information used to learn the embeddings.

## 4. Methods

### Benchmarking protocols

Here we briefly summarize our benchmarking protocols. Full details are left to Appendix B. We work with Perturb-Seq data from Nadig et al. (2024). For each cell line, perturbations are randomly split into train and test sets. We train two types of models: expression prediction (*k*-nearest neighbors) and function prediction (logistic regression). We use nested cross-validation: 5-fold cross-validation to robustly assess generalization performance, and 5-fold cross-validation for hyperparameter tuning.

### Metrics

For expression prediction, we measure predictive performance using average *ℓ*^2^ error, defined as:

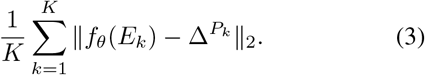

While *ℓ*^2^ error has limitations (e.g. all genes treated equally, sensitivity to outliers), we found that more complex metrics (e.g. average *ℓ*^2^ error on per-perturbation DE genes, cosine similarity, correlation) did not lead to significantly different results. For function prediction, we report macro-F1. All test metrics are computed using 5-fold cross validation.

We now describe the five categories of embeddings we benchmark. Full details can be found in Appendix C.

### Baselines

We include three *negative control* baselines to identify worst-case performance. In the **random** baseline, we generate perturbation embeddings as *E*_*k*_ *∼ U* ([0, 1]^*p*^). These perturbation embeddings carry no information about relationships between perturbations. The **no change** baseline simply predicts that the perturbed expression is identical to the control expression. The **train mean** baseline predicts the mean of all perturbed cells in the training set. The **idealized baseline** is a *positive control* baseline which has unrealistic access to ground truth information. We build the matrix of perturbed expression profiles 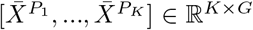 and perform PCA. This collection of embeddings reflects the ground-truth similarity between all *K* perturbations. Finally, we include a simple baseline (**PCA**) that forms gene embeddings by performing PCA on the transpose of *X*^*C*^.

### Expression-based cell FMs

We extract gene embeddings from several recent cell foundation models: AIDO.Cell (3M, 10M, and 100M parameter variants) (Ho et al., 2024), Transcriptformer (Sapiens) (Pearce et al., 2025), scPRINT (Kalfon et al., 2025), Geneformer (98M parameter variant) (Chen et al., 2024), and scGPT (Cui et al., 2024).

### DNA FMs

We use the AIDO.DNA model (Ellington et al., 2024) to compute gene embeddings based on the nucleotide sequence. For each gene, we run inference on a 4kbp window centered at the transcription start site. This yields per-nucleotide embeddings, which are then mean pooled to form gene embeddings.

### Protein FMs

We consider four models: AIDO.StructureTokenizer (Zhang et al., 2024), AIDO.ProteinIF-16B (Sun et al., 2024), ESM-2 (Lin et al., 2022), and STRING-SPACE (sequence) (Hu et al., 2024a). For AIDO.StructureTokenizer and AIDO.ProteinIF16B, we obtain an embedding for each residue and mean pool, averaging over proteins if multiple are available for a gene. For ESM-2, we obtained gene embeddings from the scPRINT API (Kalfon et al., 2025).

### Prior knowledge

We include 5 variants of embeddings from GenePT (Chen & Zou, 2024b), each trained on a different text data source. GenotypeVAE (Yu & Welch, 2022) is trained to embed genes based on the associated GO annotations. These embeddings do not vary between cell lines.

### Networks

We consider 3 network-based embeddings derived from the STRING database (Szklarczyk et al., 2025), including one from Hu et al. (2024a), STRING-SPACE (network), and 2 that we trained for this study, STRING-GNN (WaveGC) and STRING-GNN (NBFNet).

## 5. Results

### Models based on biological knowledge perform the best

In Fig. 2 we see that the best-performing models are all based on structured prior knowledge, such as descriptive text or knowledge graphs. Models based on gene expression, DNA sequences, or protein structure perform worse.

### Some cell FMs outperform simple baselines

Our results in Fig. 1 indicate that some cell FMS outperform simple mean and PCA baselines. Prior works (Kernfeld et al., 2023; Ahlmann-Eltze et al., 2024; Wenteler et al., 2024), which studied narrower collections of FMs using different protocols, did not find evidence that cell FMs could outperform simple baselines. The two cell FMs that convincingly outperform PCA (AIDO, scGPT) are the only ones that use relatively simple masked expression prediction objectives, suggesting that such objectives may be better for learning functional relationships between genes.

### AIDO.Cell shows evidence of scaling

We study variants of AIDO.Cell with different parameter counts (3M, 10M, 100M). As model size increases, we see a small but consistent improvement in performance. Model scaling may therefore be one way to improve performance for this task.

### ESM2 is the best molecular embedding

The other protein models and AIDO.DNA perform on par with random embeddings. Interestingly, AIDO.ProteinIF-16B (which learns from both protein sequence and structure information) consistently outperforms AIDO.StructureTokenizer (which learns from structure alone). We also observe that scPRINT (which uses ESM2 embeddings as inputs) performs on par with ESM2 embeddings. However, no protein embedding convincingly outperforms PCA.

### Prediction similarity is driven by modality more than model

Fig. 3 shows that embeddings learned from similar modalities tend to cluster (e.g. ESM2 and AIDO.ProteinIF16B, or the AIDO.Cell models, scGPT, and scPRINT). Exceptions include embeddings whose performance is close to random, which all cluster together regardless of modality.

### Non-cross-validated results are unreliable

The error bars in Fig. 2 show that there is significant split-to-split variability, even when we tune the hyperparameters for each split individually. This reinforces the importance of our robust evaluation protocols, without which it would be easy to draw incorrect conclusions.

### Model rankings are similar for expression prediction and function prediction

Fig. 4 shows multi-label classification results for our function prediction formulation of perturbation response modeling. Comparing with Fig. 1, which summarizes embedding performance for expression prediction, we see that the best-performing models are the same in both cases (embeddings based on biological knowledge or networks). Note that some of these embeddings were trained with GO terms, which is likely advantageous for this task. The protein models seem to perform slightly better than the expression models for function prediction, while the reverse is true for expression prediction.

## 6. Conclusion

In this study, we systematically evaluated gene embeddings derived from various families of FMs and network-based approaches on the task of perturbation response prediction. We assessed their utility both in terms of predicting gene expression changes following perturbation and in a multilabel classification task involving the prediction of perturbation-specific functional terms. Our results indicate that expression-based FMs do not yet consistently outperform more standard embedding approaches, such as those based on gene interaction networks or curated textual knowledge. Further investigation is needed to determine the biological relevance and utility of these FMs at the level of individual perturbations. See Appendix A for limitations.

## A. Limitations

By design, our study uses only one type of perturbation response model. This allows us to evaluate the effect of perturbation embeddings in isolation. We chose kNN regression for its simplicity and efficiency. It is possible that different perturbation response models exhibit different trends. However, prior work indicates that simple baselines tend to outperform more complex models (Ahlmann-Eltze et al., 2024).

In addition, our study focuses on predicting the average perturbation response of a collection of cells. Perturbation modeling can also be performed at the single-cell level (Bunne et al., 2023), and performance trends may be different in that setting.

Fine-tuning may also lead to performance improvements; we consider only fixed embeddings in this work.

Finally, we acknowledge that one model may be better than another without any of the models being “good enough” to be useful for drug discovery. The ultimate test of any perturbation response model must be wet lab validation of their predictions.

## B. Benchmarking Protocol Details

Here we describe details of our benchmarking protocol that were omitted from the main paper.

### Perturbation data

We construct our benchmarks using Perturb-Seq data from four cell lines: K562 (lymphoblast from individual with chronic myelogenous leukemia), RPE1 (retinal pigment epithelial cell from a healthy individual), Jurkat (T cell from individual with leukemia), HepG2 (epithelial cell from an individual with hepatocellular carcinoma). In particular, we use the “essential” gene knockout data from Nadig et al. (2024).

### scRNA-seq data normalization

In our problem formulation, we start with *N* control cells *X*^*C*^ ∈ ℝ^*N×G*^ and *M*_*k*_ perturbed cells 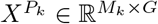 where 1 *≤ k ≤ K*. These are normalized expression matrices, not raw count matrices. For each cell, the raw counts *x*^raw^ ∈ ℝ^*G*^ are converted to normalized counts *x*^norm^ ∈ ℝ^*G*^ by computing

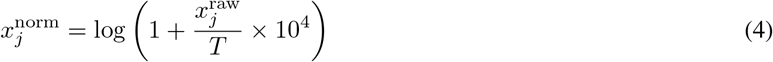

for 1 *≤ j ≤ G* where 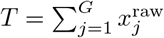 is the total count for the cell. These normalized values are used to compute average expression 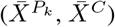 and regression targets 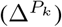.

### Splits

We split uniformly at random by perturbation. Test perturbations are never seen during training.

### Perturbation set selection

In general, each collection of perturbation embeddings can cover a different set of genes for each cell line. For each cell line, we only benchmark the perturbations (i.e. genes) that are present in all embedding collections. All other perturbations are discarded. For K562, we retain 1534 / 2053 perturbations (*∼* 75%). For the other three cell lines, we retain 1752 / 2386 perturbations (*∼* 73%).

### Embedding preprocessing

Different models produce embeddings of different dimensionality, and higher-dimensional embeddings tend to be more expressive. To control for this confounder, we compute PCA for each embedding and project to 100 dimensions. Features are standardized via *z*-scaling prior to PCA.

### Expression prediction model

For our expresion prediction formulation, we train *k*-nearest neighbor regressors as implemented in `scikit-learn`. We tune the number of neighbors *n*_*b*_ using 5-fold cross-validation. We consider *n*_*b*_ ∈ [20, 40, 60, 80, 100] and select the model with the lowest *ℓ*^2^ error. All other parameters are left at their default values.

### Function prediction model

For our function prediction formulation, we train logistic regression models as implemented in `scikit-learn`. In particular, we use `MultiOutputClassifier` with `LogisticRegression` and `n_iters` = 500. All other parameters are left at their default values.

### Gene set enrichment analysis

We restrict our analysis of function prediction to the K-562 cell line, since the ranking of different embeddings is consistent across all four lines. For each perturbation *k ∈* [*K*], we compare the expression values of 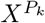 against 5000 randomly sampled control cells *X*^*C*^ using gene-wise *t*-tests. Genes with significant enrichment (FDR *≤* 0.05, Benjamini Hochberg correction (Benjamini & Hochberg, 1995)) are passed to gene set enrichment analysis (GSEA) (Subramanian et al., 2005) using gene ontology (GO) annotations (Ashburner et al., 2000), as implemented in GSEApy (Fang et al., 2023). We encode each perturbation *k* as a multilabel vector of GO terms that reach FDR *≤* 0.05. We discard terms associated with fewer than 20 perturbations, which yields a final label set of 966 GO terms for our function-prediction benchmark.

## C. Embedding Generation

This section gives additional details for some embedding generation protocols that were not fully described in the main text.

### C.1. Prior Knowledge

#### GenotypeVAE

The original GenotypeVAE was trained with a latent dimension of 10 (Yu & Welch, 2022). For the embeddings in this work, we retrained the model with a latent dimension of 128. Additionally, we used the latest gene annotations from functionome (Feuermann et al., 2025) to encode genes. All hyperparameters were identical to those from Yu & Welch (2022).

### C.2. Networks

We evaluated four network-based protein embedding strategies, grouped into two categories: (1) **STRING-SPACE** embeddings directly adopted from prior work, and (2) **STRING-GNN** embeddings developed and trained in this study using a shared framework based on STRING networks and contrastive edge learning.

### C.3. STRING-SPACE

We adopted two pretrained embeddings from the SPACE framework by Hu et al. (Hu et al., 2024b): **STRING-SPACE (sequence)** and **STRING-SPACE (network)**. Sequence-based embeddings were generated using the transformer model *ProtT5-XL-UniRef50* (Elnaggar et al., 2022), in which residue-level embeddings were extracted from full-length protein sequences and mean-pooled to produce a 1024-dimensional vector per protein. Network-based embeddings were obtained by applying weighted *node2vec* to species-specific STRING protein-protein association networks, resulting in 128-dimensional node representations. These were aligned across over 1,300 species using a modified *FedCoder* framework, which projects species-specific embeddings into a unified 512-dimensional latent space via orthology-guided autoencoders.

### C.4. STRING-GNN

We trained two GNN-based models—**STRING-GNN (WaveGC)** and **STRING-GNN (NBFNet)**—on the human STRING network (v11.5), using combined scores as edge weights and a shared contrastive training objective. In both models, input node features were initialized with the 1024-dimensional *STRING-SPACE (sequence)* embeddings, then projected to 128 dimensions using a learnable linear layer. Final node embeddings were passed into an MLP edge decoder and optimized using an InfoNCE contrastive loss over observed (positive) and sampled (negative) edges.

### STRING-GNN (WaveGC)

employs spectral graph convolution via learnable graph wavelets (Liu et al., 2024). The model applies 2 layers of WaveGC using 3 wavelet scales and 7 Chebyshev terms, enabling multi-resolution filtering that captures both local and long-range interactions. Parameters were learned using Adam optimizer (Kingma & Ba, 2014) with a learning rate of 1*e*^*−*3^ for 200 training epochs and early stopping with a tolerance of 30 epochs based on validation loss.

#### STRING-GNN (NBFNet)

uses the Neural Bellman-Ford framework (Zhu et al., 2021), which generalizes classical path-based link prediction by learning INDICATOR, MESSAGE, and AGGREGATE functions. The model performs 2 message-passing iterations to build node-pair representations, which are scored using the same MLP decoder and InfoNCE loss, with identical optimization settings as in STRING-GNN (WaveGC).

### C.5. Cell FMs

Embedding generation protocols for different cell FMs are identified in Tab. 1 and defined below. For Transcriptformer, scPRINT, Geneformer, and scGPT we compute per-gene embeddings by computing the average embedding for each gene across all control cells for each cell line. For AIDO.Cell, we compute the gene embeddings using the K562 control cells from Norman et al. (2019). This means that the AIDO.Cell embeddings are at a disadvantage for the other three cell lines.

#### Fixed gene set protocol

Some models (e.g. AIDO.Cell) have fixed gene sets – for every cell, they produce embeddings of the same set of genes. For these models, it is straightforward to generate gene embeddings. For each cell line, we pass all control cells through the model. This yields an embedding for each gene. We average these gene embeddings over all control cells.

#### Variable gene set protocol

Some models (e.g. TranscriptFormer) have variable gene sets. These models apply some sort of selection rule to the genes of each cell and provide embeddings for those genes only. For each cell line, we pass all control cells through the model. This yields a collection of (gene, embedding) pairs. We average the embeddings for each gene to produce final gene embeddings. Note that, due to the variable gene sets, some gene embeddings may be derived from more cells than others.

**Table 1.** Embedding generation protocols used for different cell FMs.

| Model | Protocol |
| --- | --- |
| AIDO.Cell 3M | Fixed |
| AIDO.Cell 10M | Fixed |
| AIDO.Cell 100M | Fixed |
| Geneformer | Variable |
| scGPT | Fixed |
| scPRINT | Fixed |
| Transcriptformer | Variable |

